# A novel method to sort and enrich sensory neurons

**DOI:** 10.1101/2025.11.19.688847

**Authors:** Zerina Kurtović, Sven David Arvidsson, Juan Antonio Vazquez Mora, Sijing Ye, Alex Bersellini Farinotti, Nils Simon, Emerson Krock, Lisbeth Haglund, Michael Hagemann-Jensen, Harald Lund, Camilla I. Svensson

**Affiliations:** Department of Physiology and Pharmacology, Karolinska Institutet, Stockholm, 171 76, Sweden; Alan Edwards Centre for Research on Pain, Faculty of Dental Medicine and Oral Health Sciences, McGill University, Montreal, QC, Canada; Shriners Hospital for Children and Department of Surgery, Orthopaedic Research Laboratory, McGill University, Montreal, QC, Canada; Department of Cell and Molecular Biology, Karolinska Institutet, Stockholm, 171 76, Sweden

## Abstract

Peripheral sensory neurons, residing in the dorsal root ganglia (DRG), relay sensory information from the periphery to the central nervous system. Although single-cell transcriptomic studies have identified over 20 distinct sensory neuron subtypes, functional analysis and assessment of subtype-specific pathological changes remain difficult. Effective isolation and enrichment of sensory neurons are challenging yet essential for functional studies. Therefore, we used single-cell transcriptomic data from DRG to identify a panel of neuronal surface markers, including *Nrxn2* and *Pirt*. Using these markers, we developed a fluorescence-activated cell sorting (FACS) panel for neuronal enrichment and analysis that does not rely on transgenic mouse strains and can be broadly applied. The panel was validated by microscopy and single-cell RNA (scRNA) sequencing, which also revealed broad representation of neuronal subtypes. Expression of these markers in human DRG underscores the translational value of this isolation method for sensory and pain studies. Overall, this study provides a valuable tool for isolating DRG neurons, advancing research on sensory neuron function and pain biology, and facilitating neuroimmune studies.

## Main

The DRG contains cell bodies of diverse peripheral sensory neuron subtypes specialized in distinct sensory modalities, extending axons to the spinal cord and peripheral tissues. In addition to sensory neurons, DRGs contain diverse non-neuronal cell types, contributing to a heterogeneous cellular environment^1^. This complexity poses substantial challenges for studying sensory neurons in isolation. Accurate molecular characterization of sensory neurons and their function requires methods that can selectively isolate these cells while maintaining a broad coverage across all neuronal subtypes. However, the lack of broadly applicable and specific surface markers for neurons has limited efficient FACS sorting of DRG neurons for downstream analyses such as cell-based transcriptomics, proteomics or flow cytometry-based assays.

Previous studies addressing neuronal isolation have used fluorescent reporter mice to sort DRG neurons, injected retrograde tracing dyes to isolate subsets of neurons based on their innervation target or used neuronal nuclei for transcriptomic studies^1–6^. These studies have revealed significant transcriptomic differences between isolated neurons and the cellularly heterogeneous whole DRG tissue, highlighting the importance of isolating neurons^2^. However, the above approaches have important limitations. Reporter-based methods depend on the availability of transgenic mouse strains, which are predominantly on the C57BL/6 background. Retrograde tracing dyes are restricted to specific neuronal subpopulations. Although nuclear isolation enables transcriptomic profiling, it is more limited than whole-cell approaches and precludes the use of powerful protein-level co-analysis methods such as CITE-seq and flow cytometric assessment of surface marker expression. As a result, the ability to analyze representative DRG neuron populations across diverse mouse strains and species in a broadly applicable, reproducible and scalable manner remains a major methodological challenge.

To overcome these challenges, we analyzed snRNA sequencing data from L3-L5 mouse DRGs to identify genes specific to neurons (Kurtović et al, in preparation). This analysis revealed distinct neuronal, glial, immune, vascular and fibroblast cell clusters. Differential gene expression analysis between neurons and non-neuronal clusters revealed 117 genes that were expressed in more than 99% of neurons and fewer than 15% of non-neuronal cells (adjusted p < 0.05). To pinpoint potential surface markers for neuronal isolation, we screened these genes for transmembrane proteins with available antibodies against extracellular epitopes, yielding 17 candidates. Among these, antibodies against NRXN2, PIRT and GABRR2 were suitable for immunostaining. We then investigated whether these proteins could serve as novel tools for isolating DRG neurons independent of genetic labeling (Fig. 1a). As the GABBR2 signal did not differ from the fluorescence minus one (FMO) control, only NRXN2 and PIRT were included in the final panel.

**Figure 1.**
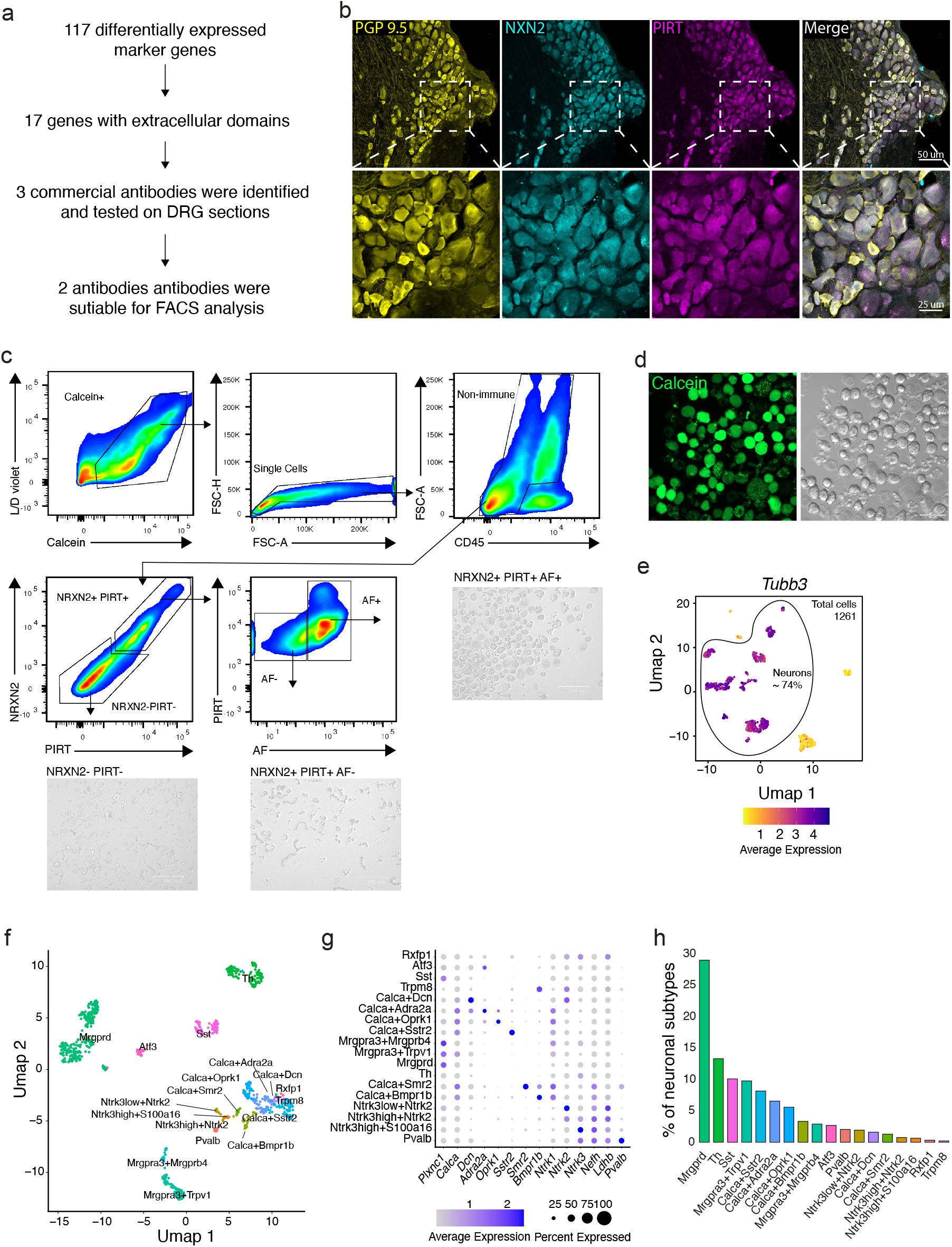
Neuronal antibody panel for isolation of sensory neurons. (a) Workflow for identification of neuronal surface markers. (b) Immunostainings of mouse DRGs for PGP9.5, NRXN2 and PIRT validates protein expression in PGP9.5+ sensory neurons. White signal in the merged image indicates co-expression. Scale bars 25-50 um. (c) Gating strategy for sorting of and images of sorted cell populations: triple negative, NRXN2+ PIRT+AF- and NRXN2+ PIRT+AF+. The images show that the target gate is enriched for sensory neurons while few neurons are outside the gate. Scale bar 100 um. (d) Confocal microscopy of sorted Calcein+ (live) NRXN2+ PIRT+AF+ neurons. Scale bar 50 um. (e) Smart-seq3xpress analysis of NRXN2+ PIRT+AF+ cells shows high *Tubb3* expression in majority of clusters (n=1261). (f) Neuronal subtypes are labelled based on SingleR annotation using mouse cells from the DRG atlas^1^ as reference (n=933). (g) Expression of marker changes confirms the SingleR labels. (h) Percentage of neuronal subtypes among all neurons in the Smart-seq3xpress analysis.

PIRT, a regulator of TRPV1 activity, is selectively expressed in DRG but not in central nervous system neurons^7^, whereas NRXN2 is present in both peripheral and central nervous systems where it contributes to synaptic function^8^. Immunostaining of DRG sections confirmed neuronal expression of both markers at the protein level, demonstrated by their co-localization with PGP9.5 (Fig. 1b). While both genes showed low-frequency RNA expression in some non-neuronal cells, we only observed rare PIRT protein expression in non-neuronal cells in tissue sections, primarily localized to the DRG capsule (Fig. 1b).

To further assess whether these markers could be used for neuronal isolation, we performed FACS sorting using the same PIRT and NRXN2 antibodies conjugated to fluorophores (Fig 1c). This revealed that a subset of NRXN2+ PIRT+ live cells showed high autofluorescence in channels excited by the 640 nm laser. We therefore collected three populations: NRXN2+ PIRT+ Autofluorescence (AF)+ cells, NRXN2+ PIRT+AF-cells and triple-negative live cells. Microscopy analysis of collected cells validated that the NRXN2+PIRT+AF+ fraction was highly enriched for neurons (Fig. 1c,d) whereas the NRXN2+ PIRT+AF- and NRXN2-PIRT-gates contained few neurons (Fig. 1c).

Next, we performed scRNA sequencing of NRXN2+PIRT+AF+ cells using the Smart-seq3xpress^9^ platform. This analysis confirmed that roughly 74% of the sequenced cells passing quality control were sensory neurons. The remaining clusters comprised multiplets, mainly of glial and endothelial origin, demonstrating an increase in the neuronal fraction from 1–4%^10^,^11^ to 74% (Fig. 1e). Cell-type annotation using SingleR^12^ and reference data from the DRG atlas^1^, revealed a broad representation of sensory neuron subtypes, with frequencies comparable to recent spatial transcriptomic datasets^13^, demonstrating the broad applicability of our isolation strategy (Fig. 1f-h).

To assess the translational relevance of this marker panel, we examined human DRG neuron data from the same atlas^1^. All neuronal subsets expressed both *NRXN2* and *PIRT*, although *NRXN2* levels were lower overall (Fig. 2a). Immunostaining of DRG sections from human donors further confirmed protein expression of NRXN2 and PIRT across DRG neurons of different soma diameters (Fig. 2b). These results suggest that this marker combination may be broadly applicable for also identifying human DRG neurons.

**Figure 2.**
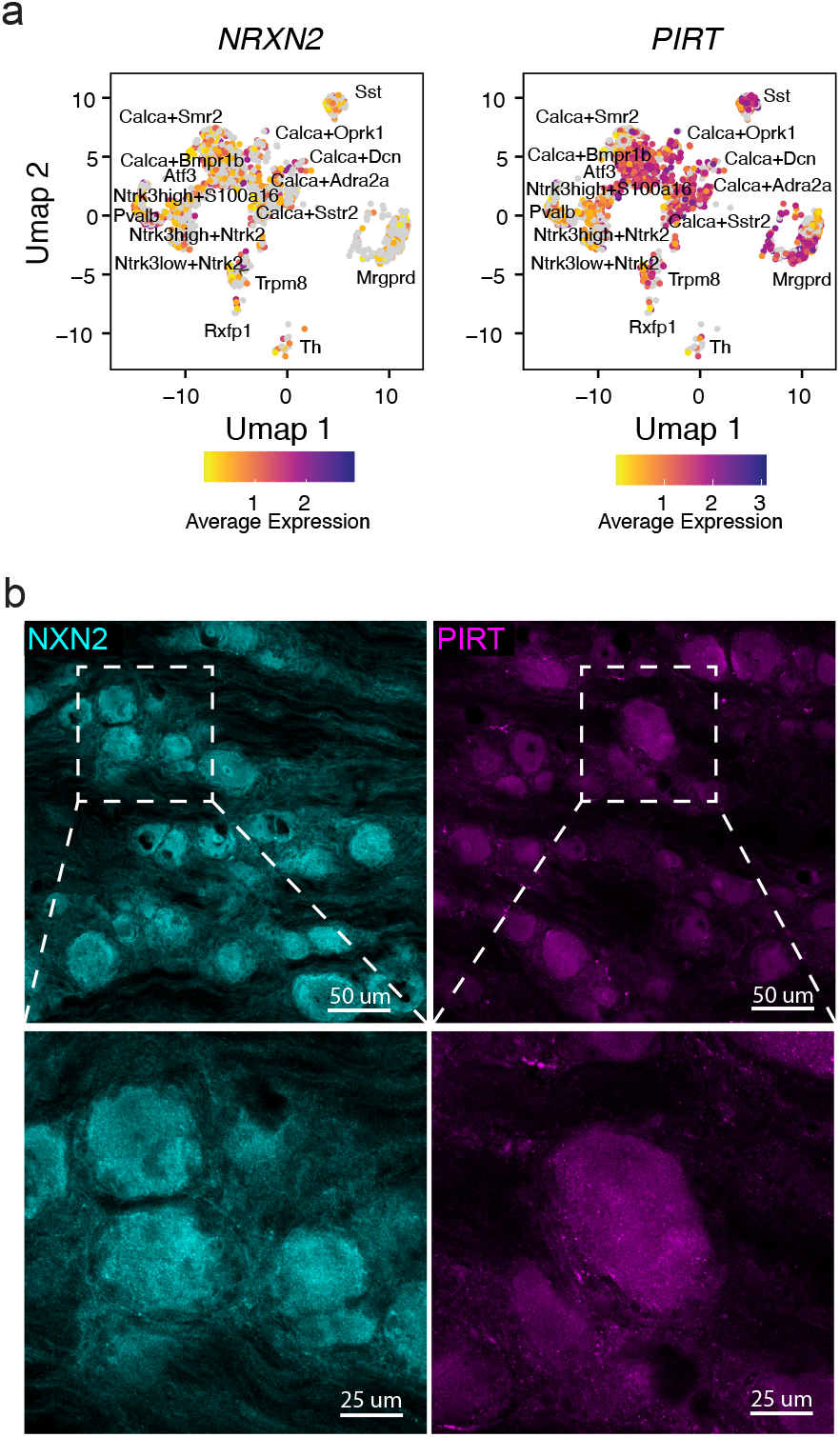
NRXN2 and PIRT label human DRG neurons. (*a*) Expression of *NRXN2* and *PIRT* in human DRG neurons from a publicly available DRG atlas^1^. (*b*) Immunostaining of *NRXN2 and* PIRT in human DRG. Scale bars 25-50 um.

Over the past decade, single-cell sequencing studies have identified 11-18 different sensory neuron types, but several subtypes are functionally uncharacterized, particularly beyond modalities described in the skin^1,14^. The ability to isolate sensory neurons will facilitate functional studies of these transcriptionally defined subtypes *in vitro*. Importantly, with the increasing availability of human DRG tissue, this tool has the potential to enable functional studies with human sensory neurons bridging the gap between transcriptomic profiling and experimental validation.

Beyond enabling enrichment of neurons in DRG single-cell datasets without relying on transgenic models, this panel also permits simultaneous analysis of sensory neurons and non-neuronal cell types, facilitating integrated studies of neuron-immune interactions, which will be a useful tool for the growing field of peripheral neuroimmunology. Additionally, it supports sensory neuron phenotyping and a broad range of flow cytometry-based molecular assays, thereby advancing the experimental toolkit for probing sensory neuron biology at a functional level.

## Methods

### Preparation of single cell suspensions

Dataset produced from other work was reanalyzed (Kurtović et al, in preparation). Briefly, the samples were prepared in the bellow described way.

Mice were euthanized through an intraperitoneal overdose of pentobarbital (338327; APL) and subsequently perfused with ice-cold PBS. Dorsal root ganglia (DRGs) were dissected and maintained in cold PBS. For 10X single-cell sequencing, bilateral L3-L5 DRGs were collected, while flow cytometry and FACS analysis involved pooling approximately 40 DRGs. Tissue digestion was performed at 37°C for 40 minutes using an enzymatic mixture containing 2 mg/ml Collagenase I (17100-017; Gibco), 5 mg/ml Dispase II (D4693; Sigma-Aldrich), and 0.5 mg/ml DNAse I (11284932001; Roche). Samples designated for scRNA sequencing were treated with 54 µg/ml actinomycin D (000025-1890, Sigma) to inhibit transcriptional changes. The resulting cell suspension was filtered through a 40-50 µm strainer, and enzymatic activity was stopped by adding 2 mM EDTA. Myelin debris was eliminated using a 38% Percoll gradient (GE17-0891-02; Sigma-Aldrich), followed by washing and resuspension of the cells in PBS.

### 10x Genomics scRNA-seq preparation and analysis

Single-cell suspensions of DRG cells from BALB/cAnNRj mice were incubated with an FcR blocking solution (No. 130-092-575; Miltenyi Biotec) to prevent nonspecific binding. Cells were then stained with Calcein Red-AM (425205; BioLegend) and the LIVE/DEAD™ Fixable Violet Dead Cell Stain (L34955, Invitrogen). Additionally, TotalSeq anti-mouse Hashtag antibodies (BioLegend) were used to label individual samples. Live, single cells (Calcein^+^, LIVE/DEAD^−^) were sorted using a BD FACSAria Fusion instrument at the Biomedicum Flow Cytometry Core Facility. Library preparation and sequencing (3’ GEM-Xv4) were conducted at the SciLifeLab sequencing facility in Solna using a NovaSeq 6000 platform, achieving 70,785 reads per cell. The resulting 10X CellRanger output files (barcodes, feature, and count matrices) were processed in R Studio using Seurat^15^. Quality control filtering retained cells with more than 700 reads and less than 5% mitochondrial content. Data normalization and scaling were performed using the Seurat functions *NormalizeData* and *ScaleData*. Highly variable features were identified using the variance-stabilizing transformation (vst) method in *FindVariableFeatures*, with the top 2,000 features selected for principal component analysis (*RunPCA*). Clustering analysis involved computing K-nearest and shared nearest-neighbor graphs using *FindNeighbors* with 26 dimensions. Graph-based clustering was performed using the Louvain algorithm (*FindClusters*) with a resolution of 0.5, informed by cluster tree analysis (*Clustree*). Novel neuronal marker genes were identified via differential gene expression (*FindAllMarkers*). Cell type annotation was based on established marker genes. Identified neuronal genes were screened for transmembrane expression and appropriate antibodies through ChatGPT-assisted manual analysis.

### FACS analysis

Single-cell suspensions of DRG cells isolated from male and female BALB/cAnNRj mice (n=5) were treated with an FcR blocking solution (No. 130-092-575; Miltenyi Biotec). The cells were subsequently stained with Calcein AM (425201; BioLegend) and LIVE/DEAD™ Fixable Violet Dead Cell Stain (L34955, Invitrogen). The following primary antibodies were used: NRXN2 (BS-11104R, ThermoFischer), PIRT (20990-1-AP, Proteintech) and CD45:BV605 (103139, BioLegend). NRXN2 and PIRT were conjugated using specific labeling kits: PE/R-Phycoerythrin Conjugation Kit - Lightning-Link (ab102918, Abcam) and APC Conjugation Kit - Lightning-Link (ab201807, Abcam) respectively. Staining was performed at 4°C for 30 minutes, after which the cells were washed and resuspended in PBS. Viable, single cells (Calcein^+^, LIVE/DEAD^−^) were then sorted using a BD FACSAria Fusion instrument. The flow cytometry sorting was performed at the Biomedicum Flow cytometry Core facility that receives funding from the Infrastructure Board at Karolinska Institutet. Flow cytometry data was analyzed using FlowJo 10 software.

### Smart-seq3xpress library preparation and sequencing

Single NRXN2+PIRT+AF+ cells were sorted into 0.3uL Smart-seq3xpress lysis buffer and processed according to previously published protocol (ref 9) with the following project specific alterfications; Preamplification was carried out using 13 cycles of PCR, followed by tagmentation of diluted cDNA using 0.005uL TDE1 (Illumina, USA) per well. Finally, the tagmented libraries were amplified using cell specific index primers for 14 cycles of PCR. Sequencing ready Smart-Seq3xpress libraries were converted to circular ssDNA libraries using the MGIEasy Universal Library Conversion kit App-A (MGI) before sequenced on MGI DNBSEQ G400 platform using universally compatible (App-D) sequencing reagents according to the manufacturer’s instructions.

### Smart-seq3xpress preprocessing and analysis

FASTQ files were processed with zUMIs (v.2.9.7). UMI containing reads were identified by finding the pattern ATTGCGCAATG allowing for two mismatches. Reads were filtered for low quality cell barcodes (4 bases < 20 Phred score) and UMIs (3 bases < 20 Phred score) and mapped to the mouse genome (mm39) using STAR (v2.7.3a). Readcounts and UMIcounts were quantified using GENCODE GRCm39.vM29 annotation. After preprocessing a quality control filter with the following parameters was applied to cells; more than 40 % read-pairs mapped to exons+introns, more than 50.000 read-pairs sequenced, less than 10% mitochrondrial reads. Furthermore, a gene was required to be expressed in at least five cells. Analysis on the quality filtered data was performed using Seurat. Data was normalized, using LogNormalize, scaled and top 3000 variable genes and 50 principle components considered for PCA, cluster analysis and UMAP projection using Louvain at resolution 0.8.

Clusters expressing the neuronal marker *Tubb3* were subseted and the PCA were recomputed with 2000 most variable genes and 50 PCs. The reclustering was done at resolution 0.5.

### Re-analysis of DRG Atlas

The DRG atlas^1^ was subsetted to contain only mouse or only human cells/nuclei in R studio and the expression of marker genes was visualized using *scCustomize*. Label transfer of neuronal subtypes to the sequenced neurons was done using SingleR.

### Immunohistochemistry of mouse tissues

Tissues were collected from three female BALB/cAnNRj control mice that underwent perfusion with 4% formaldehyde, followed by post-fixation for 4 to 24 hours before dissection. The fixation method was selected based on subsequent staining requirements. Samples were embedded in OCT (361603E, VWR) and cryosectioned at 10–25 um thickness using a cryostat (NX70, Epredia), then mounted onto SuperFrost Plus glass slides (06400X20, Epredia). For staining, sections were thawed at room temperature for 30 min, rinsed in PBS for 10 minutes, and blocked with 3% donkey serum (S30, Sigma) in PBS containing 0.2% Triton X-100 for 30 minutes. Primary antibody incubation was performed overnight at 4°C with NRXN2 (BS-11104R, Thermo Fischer Scientific), PIRT (20990-1-AP, Proteintech), and PGP9.5 (NB110-58872, Bio-Techne).NRXN2 and PIRT were conjugated using the PE/R-Phycoerythrin Conjugation Kit - Lightning-Link (ab102918, Abcam) and the APC Conjugation Kit - Lightning-Link (ab201807, Abcam), respectively. Antibodies were diluted in 0.2% Triton X-100. PGP9.5 was detected using a Cy2-conjugated donkey anti-chicken IgY (H+L) secondary antibody (703-225-155, Jackson ImmunoResearch) diluted in PBS containing 0.2% Triton X-100 and 3% donkey serum (S30, Sigma). After additional washing, sections were counterstained with DAPI and mounted using ProLong Gold (Thermo Fisher Scientific).

### Immunohistochemistry of human tissues

Two human lumbar DRG tissues were collected in Canada from one donor (Male, 61 y.o.) after obtaining consent from the next of kin via a collaboration with Transplant Quebec (IRB approval *2019-4896*). Immediately following collection, issues were either fixed for 4–6 hours or flash frozen and fixed for 3 hours in 4% paraformaldehyde. The tissue was then cryoprotected in 30% sucrose until the tissue sunk (3-5 days) at 4°C and embedded in OCT for cryosectioning at 20 μm. Tissue sections were thawed for 1 hour at room temperature, washed twice with 0.1 M phosphate-buffered saline (PBS; pH 7.35–7.45) and once with 0.01 M PBS for 5 minutes each. Sections were permeabilized at room temperature for 30 minutes in a solution containing 0.01 M PBS, 0.16% Triton X-100, 20% dimethyl sulfoxide (DMSO), and 23 g/L glycine. After permeabilization, sections were incubated in blocking solution (0.01 M PBS, 0.2% Triton X-100, 6% normal donkey serum, and 10% DMSO) for 1 hour. Primary antibodies against neurexin and PIRT were applied at room temperature in 0.01M PBS with 0.2% Tween-20, 10 mg/L heparin, 0.02% sodium azide, 5% DMSO, and 3% normal donkey serum overnight. A donkey anti-rabbit Cy3 secondary (Jackson) was applied at room temperature for 2 hours. Sections were washed and incubated with DAPI (1:10,000) for 10 minutes, followed by one 5-minute wash in washing solution, one 5-minute wash in 0.1 M PBS, and two final washes in 0.01 M PBS. To eliminate autofluorescence, TrueBlack Plus lipofuscin quencher (Biotium, cat: 23014, diluted 1:40) was applied for 30 minutes, followed by three washes (5 minutes each) in PBS. Slides were dried for 5 minutes, then mounted using Prolong Gold Antifade reagent (Invitrogen, cat: P36934).

### Confocal imaging

Representative images were taken using a Zeiss LSM800 laser-scanning confocal microscope with a 20x objective and 0.5x digital zoom. The figures show a maximum projection from a 21 z-stack image in case of tissues and 5-stack image in case of cell suspension.

## Acknowledgements

The authors would like to acknowledge the contributions of the Biomedicum Flow cytometry Core facility, financed by the Infrastructure Board at Karolinska Institutet, for providing access to cell sorting services and technical expertise.

